# Predicting the stability of homologous gene duplications in a plant RNA virus

**DOI:** 10.1101/060517

**Authors:** Anouk Willemsen, Mark P. Zwart, Pablo Higueras, Josep Sardanyés, Santiago F. Elena

## Abstract

One of the striking features of many eukaryotes is the apparent amount of redundancy in coding and non-coding elements of their genomes. Despite the possible evolutionary advantages, there are fewer examples of redundant sequences in viral genomes, particularly those with RNA genomes. The low prevalence of gene duplication in RNA viruses most likely reflects the strong selective constraints against increasing genome size. Here we investigated the stability of genetically redundant sequences and how adaptive evolution proceeds to remove them. We generated plant RNA viruses with potentially beneficial gene duplications, measured their fitness and performed experimental evolution, hereby exploring their genomic stability and evolutionary potential. We found that all gene duplication events resulted in a loss of viability or significant reductions in fitness. Moreover, upon evolving the viable viruses and analyzing their genomes, we always observed the deletion of the duplicated gene copy and maintenance of the ancestral copy. Interestingly, there were clear differences in the deletion dynamics of the duplicated gene associated with the passage duration, the size of the gene and the position for duplication. Based on the experimental data, we developed a mathematical model to characterize the stability of genetically redundant sequences, and showed that the fitness of viruses with duplications is not enough information to predict genomic stability as a recombination rate dependent on the genetic context – the duplicated gene and its position – is also required. Our results therefore demonstrate experimentally the deleterious nature of gene duplications in RNA viruses, and we identify factors that constrain the maintenance of duplicated genes.

## Introduction

Gene duplication results in genetic redundancy; in other words, the existence of genetic elements that encode for the same function. It is a powerful process that can regulate gene expression, increase the genetic and environmental robustness of organisms, and act as a stepping stone to the evolution of new biological functions. Therefore, it is not surprising that gene duplication is a frequent phenomenon in many organisms (Zhang 2003; Andersson & Hughes 2009).

There are few examples of genetic redundancy in viral genomes. In general, viral genomes tend to be highly streamlined, with limited intergenic sequences and in many cases overlapping open reading frames (ORFs), suggesting genome size is under strong selection (Lynch 2006). RNA viruses typically have smaller genomes than DNA viruses, and consequently there is an extreme low prevalence of gene duplication in RNA viruses (Belshaw et al. 2007; Belshaw et al. 2008; Simon-Loriere & Holmes 2013). For the reverse-transcribing viruses, three different gene duplication events have been reported within the *Retroviridae* family (LaPierre et al. 1999; Kambol et al. 2003; Tristem et al. 1990). This low prevalence of gene duplication in retroviruses is surprising, since repeated sequence elements of endogenous retroviruses are thought to mediate genomic rearrangements, including gene duplication (Hughes & Coffin 2001). For the ss(-)RNA viruses, two different tandem gene duplications have been reported (Walker et al. 1992; Blasdell et al. 2012; Gubala et al. 2010; Simon-Loriere & Holmes 2013) within the *Rhabdoviridae* (infecting vertebrates, invertebrates and plants). For the ss(+)RNA viruses, single duplication events have been reported for three different domains: (*i*) a tandem duplication of the coat protein gene (CP) within the *Closteroviridae* (infecting plants) (Boyko et al. 1992; Fazeli & Rezaian 2000; Tzanetakis et al. 2005; Tzanetakis & Martin 2007; Kreuze et al. 2002; Simon-Loriere & Holmes 2013); (*ii*) a tandem duplication of the genome-linked protein gene (*VPg)* in *Foot-and-mouth disease virus* from the *Picornaviridae* (infecting vertebrates) (Forss & Schaller 1982); and (*iii*) a duplication of the third segment, generating an additional one, in *Beet necrotic yellow vein virus* from the *Benyviridae* (Simon-Loriere & Holmes 2013). To date, no cases of gene duplication in dsRNA viruses have been reported.

The variation in genome size and structure indicates that gene duplication must have played a role in the early diversification of virus genomes. However, the rapid evolution of RNA viruses and the potential fitness costs associated with harboring additional genetic material probably makes it unlikely to detect viruses with duplications, or even the signatures of recent duplication events. Strong selective constraints against increasing genome sizes are thought to play a role in the lack of gene duplications that we nowadays observe in RNA viruses (Holmes 2003; Belshaw et al. 2007; Belshaw et al. 2008). One of these constraints is the high mutation rates of RNA viruses, which is approximately one mutation per genome and per replication event (Sanjuán et al. 2010). This limits the probability of copying without errors a genome above the length limit imposed by Eigen’s error threshold (Eigen 1971): the inverse of the per site mutation rate. Another constraint is the need for fast replication due to strong within-cell and within-host competition (Turner & Chao 1998). An increase in genome size is therefore likely to increase the number of deleterious mutations that occur per genome during each round of replication, and to slow down the replication process. On the other hand, the small and streamlined RNA virus genomes also limit sequence space for the evolution of novel functions, and in turn adaptation to environmental changes.

Here, we therefore consider experimentally the evolutionary fate of gene duplications in viral genomes, in terms of their effects on fitness, the stability of the duplicated gene and the evolvability of these viruses. We experimentally explore four cases of homologous duplication of genes within the *Tobacco etch virus* (TEV; genus *Potyvirus*, family *Potyviridae)* genome (Revers & Gartia 2015): (i) the multi-functional protein (HC-Pro) involved in aphid transmission, polyprotein cleavage, genome amplification, and suppression of RNA silencing, (*ii*) the main viral protease (NIa-Pro), (*iii*) the viral RNA-dependent RNA polymerase (NIb), and (*iv*) the coat protein (CP). Potyviruses are a particularly interesting system for studying the evolution of gene duplications, as they encode a single polyprotein that is auto catalytically processed into the mature gene products. For each complete (+)RNA, as well as frame-shifted transcripts for which translation terminates at P3-PIPO, there will be isostoichiometric expression of all genes. Assuming there are no unknown mechanisms that regulate gene expression, the scope for the regulation of gene expression in potyviruses could therefore be very limited. Gene duplication may represent a way to bypass these constraints and achieve higher expression of specific genes.

We speculated that the duplication of these four proteins might have widely different impacts on TEV fitness. As HC-Pro is a multifunctional protein, two copies of HC-Pro could lead to specific improvement of one or more of its functions. This potential improvement could possibly be caused by two mechanisms. Firstly, by simply producing more protein there could be an immediate benefit for one of HC-Pro’s functions. Secondly, there could be improvement of protein function when the duplicated virus is evolved, because the two gene copies can diverge and specialize on different functions. Higher levels of NIa-Pro may result in a more efficient processing of the polyprotein, making more mature viral proteins available faster for the replication process. As potyviruses have only a limited number of post-translational mechanisms for regulating gene expression levels, we predicted that the overproduction of NIa-Pro will alter the equilibrium concentrations of all the different mature peptides and thus have a major impact in TEV fitness. Higher levels of NIb may result in higher levels of transcription and faster replication of the virus and this could lead to higher levels of genome accumulation and potentially the within-host spread of infection by a greater number of virions. The cellular multiplicity of infection (MOI), which has been estimated to be as low as 1.14 virions per infected cell for TEV (Tromas et al. 2014a), might even increase. Higher levels of CP expression could allow for the encapsidation of more genomic RNA molecules without affecting the accumulation of all other mature peptides. However, in all these cases completion of the infectious cycle would still depend on the cytoplasmic amount of other limiting viral (*e.g.*, P1, P3, CI, and VPg) or host proteins.

The duplication events that we explore here could therefore conceivably have beneficial effects on TEV replication, perhaps offsetting the costs inherent to a larger genome and thereby increasing overall fitness. Moreover, especially in the case of HC-Pro, they could perhaps lead to the evolution of greatly improved or novel functions. However, given the scarcity of gene duplications in RNA viruses, we expected that the fitness costs of duplication are likely to be high, and that one of the two gene copies would be rapidly lost. If further mutations could potentially help accommodate the duplicated gene, then this could lead to interesting evolutionary dynamics: will the duplicated gene be lost or will beneficial mutations that lead to stable maintenance of the gene occur first? (Zwart et al. 2014) Moreover, as they could potentially disrupt correct processing of the polyprotein, the possibility that some of the duplications would not be viable in the first place could also not be discounted (Majer et al. 2014). To address these issues we have constructed four viruses with gene duplications and tested their viability. We subsequently evolved these viruses and determined the stability of the duplicated gene, as well as looking for signals of accommodation of the duplicated gene. Finally, we built a mathematical model to estimate key parameters from the experimental data, such as the recombination rates responsible for the deletion of duplicated genes, and to explore the evolutionary dynamics and stability conditions of the system.

## Materials and Methods

### Viral constructs, virus stocks and plant infections

The TEV genome used to generate the virus constructs, was originally isolated from *Nicotiana tabacum* plants (Carrington et al. 1993). In this study five different variants of TEV were used containing single gene duplications. Two of these virus variants, TEV-NIb_1_-NIb_9_ and TEV-NIb_2_-NIb_9_ were generated in a previous study (Willemsen et al. 2016). The other three variants were generated in this study: TEV-HCPro2-HCPro3, TEV-NIaPro_2_-NIaPro_8_ and TEV-CP_10_-CP_11_.

TEV-HCPro_2_-HCPro_3_, TEV-NIaPro_2_-NIaPro_8_ and TEV-CP_10_-CP_11_ were generated from cDNA clones constructed using plasmid pMTEVa, which consists of a TEV infectious cDNA (accession: DQ986288, including two silent mutations, G273A and A1119G) flanked by SP6 phage RNA promoter derived from pTEV7DA (GenBank: DQ986288). pMTEVa contains a minimal transcription cassette to ensure a high plasmid stability (Bedoya & Daròs 2010). The clones were constructed using standard molecular biology techniques, including PCR amplification of cDNAs with the high-fidelity Phusion DNA polymerase (Thermo Scientific), DNA digestion with *Eco31I* (Thermo Scientific) for assembly of DNA fragments (Engler et al. 2009), DNA ligation with T4 DNA ligase (Thermo Scientific) and transformation of *E. coli* DH5α by electroporation. Sanger sequencing confirmed the sequences of the resulting plasmids.

The plasmids of TEV-HCPro_2_-HCPro_3_, TEV-NIaPro_2_-NIaPro_8_ and TEV-CP10-CP11 were linearized by digestion with *Bgl*II prior to *in vitro* RNA synthesis using the mMESSAGE mMACHINE^®^ SP6 Transciption Kit (Ambion), as described in Carrasco et al. (2007). The third true leaf of 4-week-old *N. tabacum* L. cv Xanthi *NN* plants was mechanically inoculated with varying amounts (5 μg - 30 μg) of transcribed RNA. All symptomatic tissue was collected 7 dpi (days post inoculation) and stored at –80 °C as stock tissue.

### Serial passages

For the serial passage experiments, 500 mg homogenized stock tissue was ground into fine powder using liquid nitrogen and a mortar, and resuspended in 500 μl phosphate buffer (50 mM KH_2_PO_4_, pH 7.0, 3% polyethylene glycol 6000). From this mixture, 20 μl were then mechanically inoculated on the third true leaf of 4-week old *N. tabacum* plants. At least five independent replicates were performed for each virus variant. At the end of the designated passage duration (3 or 9 weeks) all leaves above the inoculated leaf were collected and stored at −80 °C. For subsequent passages the frozen tissue was homogenized and a sample of the homogenized tissue was ground and resuspended with an equal amount of phosphate buffer (Zwart et al. 2014). Then, new *N. tabacum* plants were mechanically inoculated as described above. The plants were kept in a BSL-2 greenhouse at 24 °C with 16 h light.

### Reverse transcription polymerase chain reaction (RT-PCR)

The wild-type TEV produces characteristic symptoms in the host plant. However, some of the altered genotypes show few or no symptoms and virus infection had to be confirmed by RT-PCR. To confirm infection and to determine the stability of the duplicated genes, RNA was extracted from 100 mg homogenized infected tissue using the InviTrap Spin Plant RNA Mini Kit (Stratec Molecular). Reverse transcription (RT) was performed using M-MuLV reverse transcriptase (Thermo Scientific) and the reverse primer 5‘-CGCACTACATAGGAGAATTAG-3’ located in the 3‘UTR of the TEV genome. PCR was then performed with *Taq* DNA polymerase (Roche) and primers flanking the region containing the duplicated gene copy (supplementary Table S1, Supplementary Material online). To test whether the ancestral gene copy was intact this region was also amplified for TEV-NIaPro_2_-NIaPro_8_, TEV-NIb_1_-NIb_9_ and TEV-NIb_2_-NIb_9_ viruses, where the duplicated genes are not located in tandem (supplementary Table S1, Supplementary Material online). PCR products were resolved by electrophoresis on 1% agarose gels. For those virus populations in which we detected deletions during the evolution experiment, we estimated the genome size based on the amplicon size and the genome size of the ancestral viruses.

### Fitness assays

The genome equivalents per 100 mg of tissue of the ancestral virus stocks and all evolved lineages were determined for subsequent fitness assays. The InviTrap Spin Plant RNA Mini Kit (Stratec Molecular) was used to isolate total RNA from 100 mg homogenized infected tissue. Real-time quantitative RT-PCR (RT-qPCR) was performed using the One Step SYBR PrimeScript RT-PCR Kit II (Takara), in accordance with manufacturer instructions, in a StepOnePlus Real-Time PCR System (Applied Biosystems). Specific primers for the *CP* gene were used; forward 5‘-TTGGTCTTGATGGCAACGTG-3’ and reverse 5‘-TGTGCCGTTCAGTGTCTTCCT-3’. The StepOne Software v.2.2.2 (Applied Biosystems) was used to analyze the data. The concentration of genome equivalents per 100 mg of tissue was then normalized to that of the sample with the lowest concentration, using phosphate buffer.

For the accumulation assays, 4-week-old *N. tabacum* plants were inoculated with 50 pl of the normalized dilutions of ground tissue. For each ancestral and evolved lineage, at least three independent plant replicates were used. Leaf tissue was harvested 7 dpi. Total RNA was extracted from 100 mg of homogenized tissue. Virus accumulation was then determined by means of RT-qPCR for the *CP* gene of the ancestral and the evolved lineages. For each of the harvested plants, at least three technical replicates were used in the RT-qPCR.

To measure within-host competitive fitness, we used TEV carrying an enhanced green fluorescent protein (TEV-eGFP) (Bedoya & Daròs 2010) as a common competitor. TEV-eGFP has proven to be stable up to six weeks (using 1- and 3-week serial passages) in *N. tabacum* (Zwart et al. 2014), and is therefore not subjected to appreciable eGFP loss during our 1-week long competition experiments. All ancestral and evolved viral lineages were again normalized to the sample with the lowest concentration, and 1:1 mixtures of viral genome equivalents were made with TEV-eGFP. The mixture was mechanically inoculated on the same species of host plant on which it had been evolved, using three independent plant replicates per viral lineage. The plant leaves were collected at 7 dpi, and stored at –80 °C. Total RNA was extracted from 100 mg homogenized tissue. RT-qPCR for the *CP* gene was used to determine total viral accumulation, and independent RT-qPCR reactions were also performed for the *eGFP* sequence using primers forward 5‘-CGACAACCACTACCTGAGCA-3′ and reverse 5 ‘-GAACTCCAGCAGGACCATGT-3′. The ratio (*R*) of the evolved and ancestral lineages to TEV-eGFP is then *R* = (*n_cp_ — n_eGFP_*)*/n_eGFP_*, where *n_CP_* and *n_eGFP_* are the RT-qPCR measured copy numbers of *CP* and *eGFP*, respectively. Within-host competitive fitness can then be estimated as 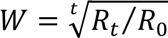 where *R*_0_ is the ratio at the start of the experiment and *R_t_* the ratio after *t* days of competition (Carrasco et al. 2007). Note that the method for determining *R* only works well when the frequency of the common is below ~0.75. This limitation was not problematic though, since in these experiments the fitness of the evolved virus populations remained the same or increased. The statistical analyses comparing the fitness between lineages were performed using R v.3.2.2 (R Core Team 2014) and IBM SPSS Statistics version 23.

### Sanger sequencing

For those evolved virus populations in which deletions were detected by RT-PCR, the exact positions of these deletions were determined. The genomes were partly sequenced by the Sanger method. RT was performed using AccuScript Hi-Fi(Agilent Technologies) reverse transcriptase and a reverse primer outside the region to be PCR-amplified for sequencing (supplementary Table S2, Supplementary Material online). PCR was then performed with Phusion DNA polymerase (Thermo Scientific) and primers flanking the deletions (supplementary Table S1, Supplementary Material online). Sanger sequencing was performed at GenoScreen (Lille, France:www.genoscreen.com; last accessed April 10, 2016) with an ABI3730XL DNA analyzer. For TEV-HCPro_2_-HCPro_3_, six sequencing reactions were done per lineage using the same outer reverse primer as used for PCR amplification plus five inner primers (supplementary Table S2, Supplementary Material online). For TEV-NIaPro_2_-NIaPro_8_, three sequencing reactions were done per lineage using three inner primers (supplementary Table S2, Supplementary Material online). For TEV-NIb_1_-NIb_9_ and TEV-NIb_2_-NIb_9_, six sequencing reactions were done per lineage using the same two outer primers as used for PCR amplification plus four inner primers (supplementary Table S2, Supplementary Material online). For TEV-CP_10_-CP_11_, two sequencing reactions were done per lineage using the same two outer primers as used for PCR amplification (supplementary Table S2, Supplementary Material online). Sequences were assembled using Geneious v.8.0.3 (www.geneious.com; last accessed April 10, 2016) and the start and end positions of the deletions were determined. Based on the ancestral reference sequences, new reference sequences were constructed containing the majority deletion variant for each of the evolved lineages.

### Illumina sequencing, variants, and SNP calling

For Illumina next-generation sequencing (NGS) of the evolved and ancestral lineages, the viral genomes were amplified by RT-PCR using AccuScript Hi-Fi(Agilent Technologies) reverse transcriptase and Phusion DNA polymerase (Thermo Scientific), with six independent replicates that were pooled. Each virus was amplified using three primer sets generating three amplicons of similar size (set 1: 5‘-GCAATCAAGCATTCTACTTCTATTGCAGC-3’ and 5‘-TATGGAAGTCCTGTGGATTTTCCAGATCC-3’; set 2: 5‘-TTGACGCTGAGCGGAGTGATGG-3’ and 5‘-AATGCTTCCAGAATATGCC-3’; set 3: 5‘-TCATTACAAACAAGCACTTG-3’ and 5‘-CGCACTACATAGGAGAATTAG-3’). Equimolar mixtures of the three PCR products were made. Sequencing was performed at GenoScreen. Illumina HiSeq2500 2×100bp paired-end libraries with dual-index adaptors were prepared along with an internal PhiX control. Libraries were prepared using the Nextera XT DNA Library Preparation Kit (Illumina Inc.). Sequencing quality control was performed by GenoScreen, based on PhiX error rate and Q30 values.

Read artifact filtering and quality trimming (3’ minimum Q28 and minimum read length of 50 bp) was done using FASTX-Toolkit v.0.0.14(hannonlab.cshl.edu/fastxtoolkit/index.html, last accessed April 10, 2016). Dereplication of the reads and 5’ quality trimming requiring a minimum of Q28 was done using PRINSEQ-lite v.0.20.4 (Schmieder & Edwards 2011). Reads containing undefined nucleotides (N) were discarded. As an initial mapping step, the evolved sequences were mapped using Bowtie v.2.2.6 (Langmead & Salzberg 2012) against their corresponding ancestral sequence: TEV (GenBank accession number KX137149), TEV-HCPro_2_-HCPro_3_ ancestral (GenBank accession number KX137150), TEV-NIaPro_2_-NIaPro_8_ ancestral (GenBank accession number KX137151), TEV-NIb_1_-NIb_9_ ancestral (GenBank accession number KT203712), TEV-NIb_2_-NIb_9_ ancestral (GenBank accession number KT203713), and against the evolved lineages including the corresponding deletions in the lineages where they are present. Subsequently, mutations were detected using SAMtools’ mpileup (Li et al. 2009) in the evolved lineages as compared to their ancestral lineage. At this point, we were only interested in mutations at a frequency > 10%. Therefore, we present frequencies as reported by SAMtools, which has a low sensitivity for detecting low-frequency variants (Spencer et al. 2014).

After the initial pre-mapping step, error correction was done using Polisher v2.0.8 (available for academic use from the Joint Genome Institute) and consensus sequences were defined for every lineage. Subsequently, the cleaned reads were remapped using Bowtie v.2.2.6 against the corresponding consensus sequence for every lineage. For each new consensus, Single nucleotide polymorphisms (SNPs) within each virus population were identified using SAMtools’ mpileup and VarScan v.2.3.9 (Koboldt et al. 2012). For SNP calling maximum coverage was set to 40000 and SNPs with a frequency < 1% were discarded.

### Modeling the stability of gene insertions

We developed a mathematical model to fit with the experimental data for the 3-week and 9-week passages. We were particularly interested in better understanding the frequency of virus populations in which viruses with deletions in the duplicated gene were present or had been fixed. The model consists of two coupled ordinary differential equations:

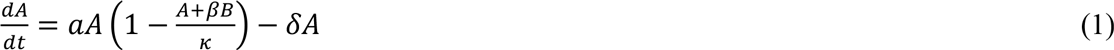

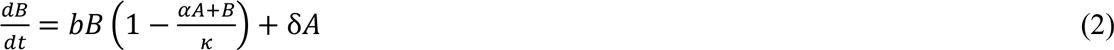

where *A* is the number of virions containing a gene duplication, *B* is the number of virions with a reversion to a single copy, *a* is the initial growth rate of *A*, *b* is the initial growth rate of *B*, *β = b/a* is a constant for determining the effect of the presence of *B* on replication of *A* (Solé et al. 1998), *α* = 1/*β* is a constant for determining the opposing effect of *A* on *B*, *κ* is the time-dependent carrying capacity of the host plant, and *δ* is the recombination rate per genome and replication at which the extra copy of the gene is removed from *A* to produce *B*. We assume that *κ* increases linearly over time, being proportional to the estimated weight of collected plant tissue (2 g for the whole plant at inoculation, 200 g for the collected leaves at 9 weeks). At the start of each round of infection, there is a fixed bottleneck size of *λ*. The number of infecting virions of *A* is determined by a random draw from a Binomial distribution with a probability of success *λA*/(*A + B*) and a size *λ*, and the number of infecting virions of *B* is then *λ* minus this realization from the Binomial distribution.

Estimates of most model parameters could be obtained from previous studies (Table 1). An estimate of *b* has been made (Zwart et al. 2012), whilst *a* can be determined knowing the competitive fitness of the virus with duplication relative to the wild-type virus. The value of *b* used is 1.344, and *a* values are 1.175, 1.234, 1.185 and 1.134 for TEV-HCPro_2_-HCPro3, TEV-NIaPro_2_-NIaPro_8_, TEV-NIb_1_-NIb_9_, and TEV-NIb2-NIb_9_, respectively. The only model parameter that needed to be estimated from the data is δ. To obtain an estimate of this parameter, we implemented the model as described in equations 1 and 2 in R 3.1.0. For each dataset to which we wanted to fit the model, we first simply ran the model for a wide range of recombination rates: considering all values of log(δ) between -20 and -0.1, with intervals of 0.1. One-thousand simulations were run for each parameter value.

**Table 1.**
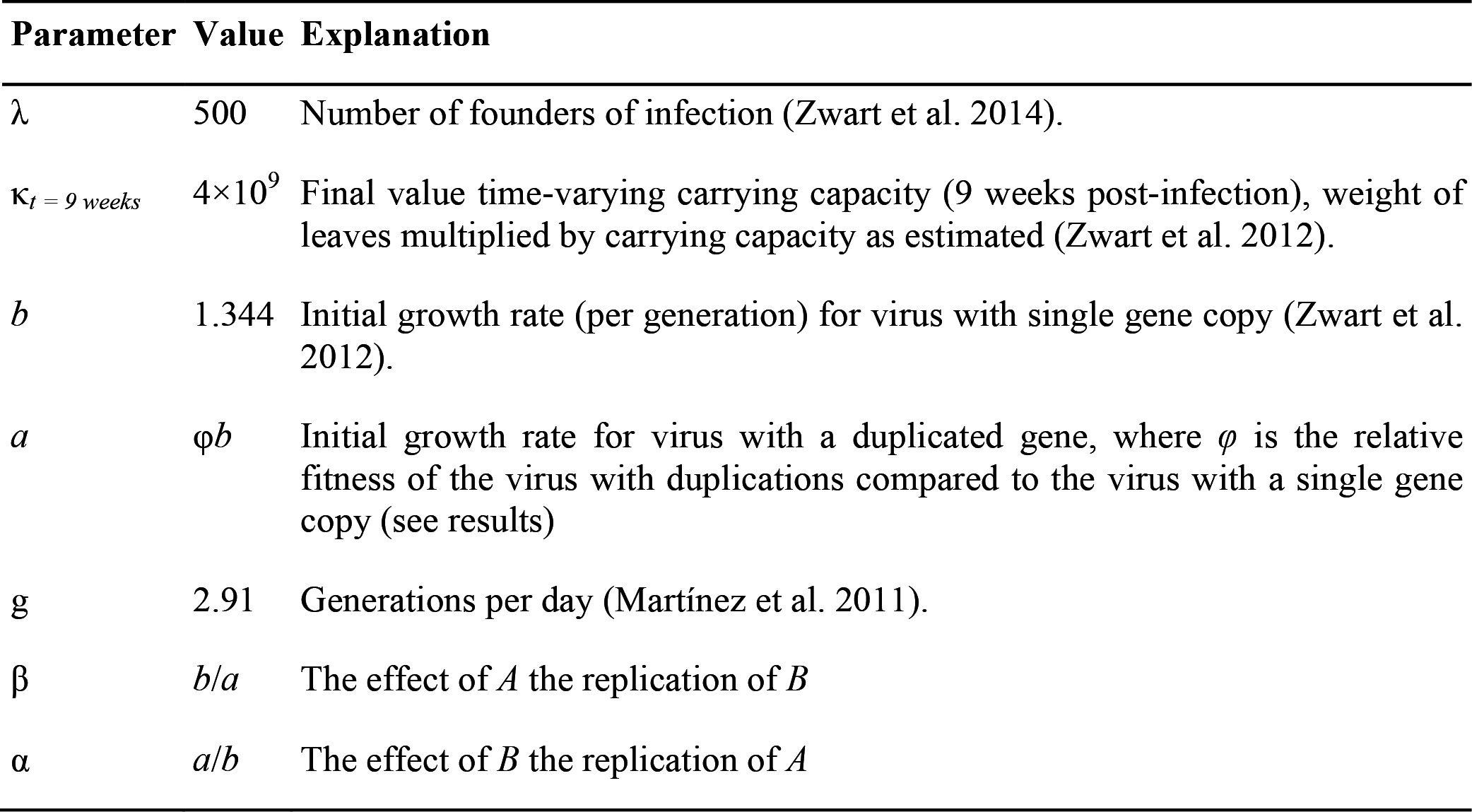
Model parameters

To fit the model to the data, we considered model predictions of the frequency of three kinds of virus populations over time: (*i*) those populations containing only the full-length ancestral virus with a gene duplication (*X*_1_), (*ii)* those populations containing only variants with a genomic deletion removing the artificially introduced second gene copy (*X*_2_), and (*iii)* those populations containing a mixture of both variants (*X*_3_). Due to the modeling of recombination and selection as deterministic processes, the model predicts recombinants will be ubiquitous. However, in order to be reach appreciable frequencies and eventually be fixed, recombinants must reach a high enough frequency that they will be sampled during the genetic bottleneck at the start of infection. Moreover, we used a PCR-based method with inherent limits to its sensitivity to characterize experimental populations. For these two reasons, we assumed that the predicted frequency of *A* must be greater than 0.1 and less than 0.9 to be considered a mixture. We then compared model predictions for the frequency of the three different kinds of virus populations with the data by means of multinomial likelihoods. The likelihood of the number of occurrences of these three stochastic variables denoting observations of a particular kind of population (*X*_1_, *X*_2_, *X*_3_) follows a Multinomial distribution with probabilities *p_1_, p_2_* and *p_3_* 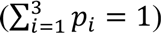. The multinomial probability of a particular realization (*x*_1_, *x*_2_, *x*_3_) is given by:

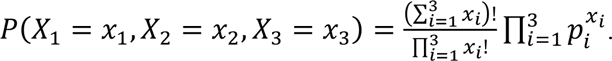

The estimate of *δ* is then simply that value that corresponds to the lowest negative log-likelihood (NLL), for the entire range of *δ* values tested. We first fitted the model with a single value of *δ* to all the data (Model 1; 1 parameter). Next, we fitted the model with a virus-dependent value of 6, but one which is independent of passage duration (Model 2; 4 parameters). We then fitted the model with *δ* value dependent on passage duration, but the same for each virus (Model 3; 2 parameters). Finally, we fit the model to each experimental treatment separately (Model 4; 8 parameters). For all these different model fittings, 95% fiducial estimates of *δ* were obtained by fitting the model to 1000 bootstrapped datasets.

The numerical solutions of the differential equations 1 and 2 used to characterize the dynamical properties, build the phase portraits and to obtain the transient times (supplementary Text S1, Supplementary Material online) have been obtained using a fourth-order Runge-Kutta method with a time step size 0.1.

## Results

### Genetic redundant constructs and the viability of the resulting viruses

To simulate the occurrence of duplication events within the TEV genome (Figure 1A), different TEV genotypes were constructed using four genes of interest (Figure 1). Each of these genotypes therefore represents a single gene duplication event. Where necessary, the termini of the duplicated gene copies were adjusted, such that the proteins can be properly translated and processed. Cleavage sites are provided, similar to the original proteolytic cleavage sites at the corresponding positions. A description of every duplication event will be given in the same order as these genes occur within the TEV genome.

**Figure 1.**
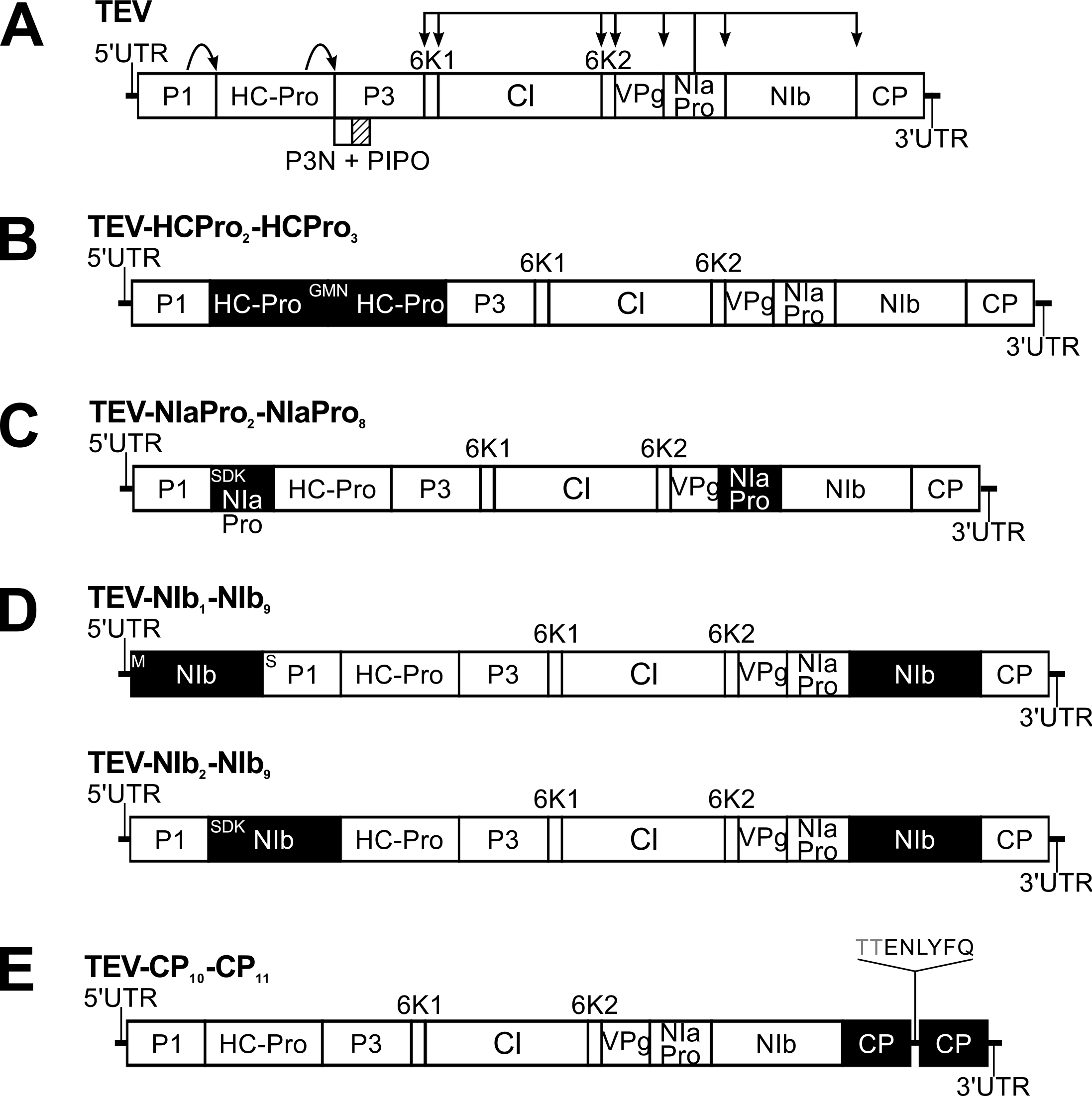
Schematic representation of the different TEV genotypes containing gene duplications. The wild-type TEV (A) codes for 11 mature peptides, including P3N-PIPO embedded within the P3 protein at a +2 frameshift. Five different viral genotypes containing a single gene duplication were constructed. Second copies of *HC-Pro* (B), *NIa-Pro* (C) and *NIb (D)* were introduced between *P1* and *HC-Pro*. A second copy of *NIb* was also introduced before *P1* (D). And a second copy of *CP* was introduced between *NIb* and *CP* (E). For simplification P3N-PIPO is only drawn at the wild-type TEV.

First, for duplication of the multifunctional *HC-Pro* gene, a second copy of *HC-Pro* was inserted in the second position within the TEV genome, between the *P1* serine protease gene and the original *HC-Pro* copy, generating a tandem duplication (Figure 1B). This position is a common site for the cloning of heterologous genes (Zwart et al. 2011). Second, a copy of the *NIa-Pro* main viral protease gene was introduced between *P1* and *HC-Pro* (Figure 1C). Third, two genotypes containing a duplication of the *NIb* replicase gene were generated (Willemsen et al. 2016), where a copy of the *NIb* gene was inserted at the first position (before *P1*) and the second position in the TEV genome (Figure 1D). Fourth, for duplication of the *CP* we introduced a second copy at the tenth position between *NIb* and *CP*, generating a tandem duplication (Figure 1E). Henceforth we refer to these five genetic redundant viruses as TEV-HCPro_2_-HCPro_3_, TEV-NIaPro_2_-NIaPro_8_, TEV-NIb_r_NIb_9_, TEV-NIb_2_-NIb_9_, and TEV-CP_10_-CP_11_, respectively, with subscripts denoting the intergenic positions of the duplicated gene in question.

The viability of these viruses was tested in *N. tabacum* plants, by inoculating plants with approximately 5 pg *in vitro* generated transcripts. TEV-HCPro_2_-HCPro_3_, TEV-NIaPro_2_-NIaPro_8_, TEV-NIb_1_-NIb_9_, and TEV-NIb_2_-NIb_9_, were found to infect *N. tabacum* plants, as determined by RT-PCR on total RNA extracted from these plants. After performing multiple viability tests, TEV-CP_10_-CP_11_ demonstrated to have a very low infectivity and high amounts (> 20 μg) of RNA are needed for infection to occur. Performing RT-PCR of the region containing two *CP* copies, we detected either (*i*) a band corresponding to the wild-type virus, indicating that upon infection with RNA the second *CP* copy is deleted immediately, or (*ii*) a band that indicates the two *CP* copies are present. Taking the latter as a starting population for experimental evolution, within the first passage, we detect a band corresponding to the wild-type virus, in six out of eight lineages, and we did not detect any infection in the remaining two lineages. When sequencing the region containing the deletions in the different lineages, using Sanger technology, exact deletions of the second *CP* copy were observed. We discontinued further experiments on this virus due to the extreme instability of the second *CP* copy.

### Evolution of genetically redundant viruses

After reconstitution of TEV-HCPro_2_-HCPro_3_, TEV-NIaPro_2_-NIaPro_8_, TEV-NIb_1_-NIb_9_, and TEV-NIb2-NIb9 from infectious clones, these viruses containing gene duplications were evolved in *N. tabacum* plants. All viruses were evolved for a total of 27 weeks, using nine 3-week passages and three 9-week passages with at least five independent lineages for each passage duration. In the starting population of TEV-HCPro_2_-HCPro_3_ we observed mild symptoms. However, in lineages from the first 3-and 9-week passages, the plants rapidly became as symptomatic as those infected by the wild-type virus. At the start of the evolution experiment, TEV-NIaPro2-NIaPro8 also displayed only mild symptoms and infection appeared to expand slower than for the wild-type TEV. However, in the first 9-week passage symptoms became stronger, similar to the wild-type virus, as the virus expanded through the plant. These stronger symptoms were also observed in the subsequent 9-week passages. Increases in symptom severity were also observed for the TEV-NIb_1_-NIb_9_ and TEV-NIb_2_-NIb_9_ viruses (Willemsen et al. 2016).

Partial and complete deletions of the duplicated gene copy were detected by RT-PCR (Figure 2), but never in the ancestral gene. Deletion of the duplicated gene copy of the TEV-HCPro_2_-HCPro_3_ variant occurred rapidly after infection of the plants; after one passage the gene duplication could no longer be detected by RT-PCR (Figure 2A). No deletions were detected in the TEV-NIaPro2-NIaPro8 lineages using the shorter 3-week passages (Figure 2B). Deletions were not detected in the first 9-week passage either, but in the second passage partial or complete deletions did occur. Mixed populations that contain virions with a deletion together with virions that have maintained the intact duplicated copy, are mainly present in the TEV-NIb_1_-NIb_9_ and TEV-NIb_2_-NIb_9_ lineages (Figures 2C and 2D). However, TEV-NIb_1_-NIb_9_ loses the duplicated copy faster (Figure 2C; second 3-week passage, and first 9-week passage) than TEV-NIb_2_-NIb_9_ (Figure 2D; third 3-week passage, and second 9-week passage). Based on the majority deletion variants observed by RT-PCR, genome size was estimated for every passage (Figure 3). Comparing the genome size of the different viral genotypes in Figure 3, there are clear differences in the time until the duplicated gene copy is deleted. The duplicated *HC-Pro* copy appears the least stable, while the duplicated *NIa-Pro* copy appears to be the most stable. For TEV-NIaPro_2_-NIaPro_8_, TEV-NIb_1_-NIb_9_, and TEV-NIb_2_-NIb_9_, there are lineages that contain deletions that lead to a genome size smaller than that of the wild-type TEV.

**Figure 2.**
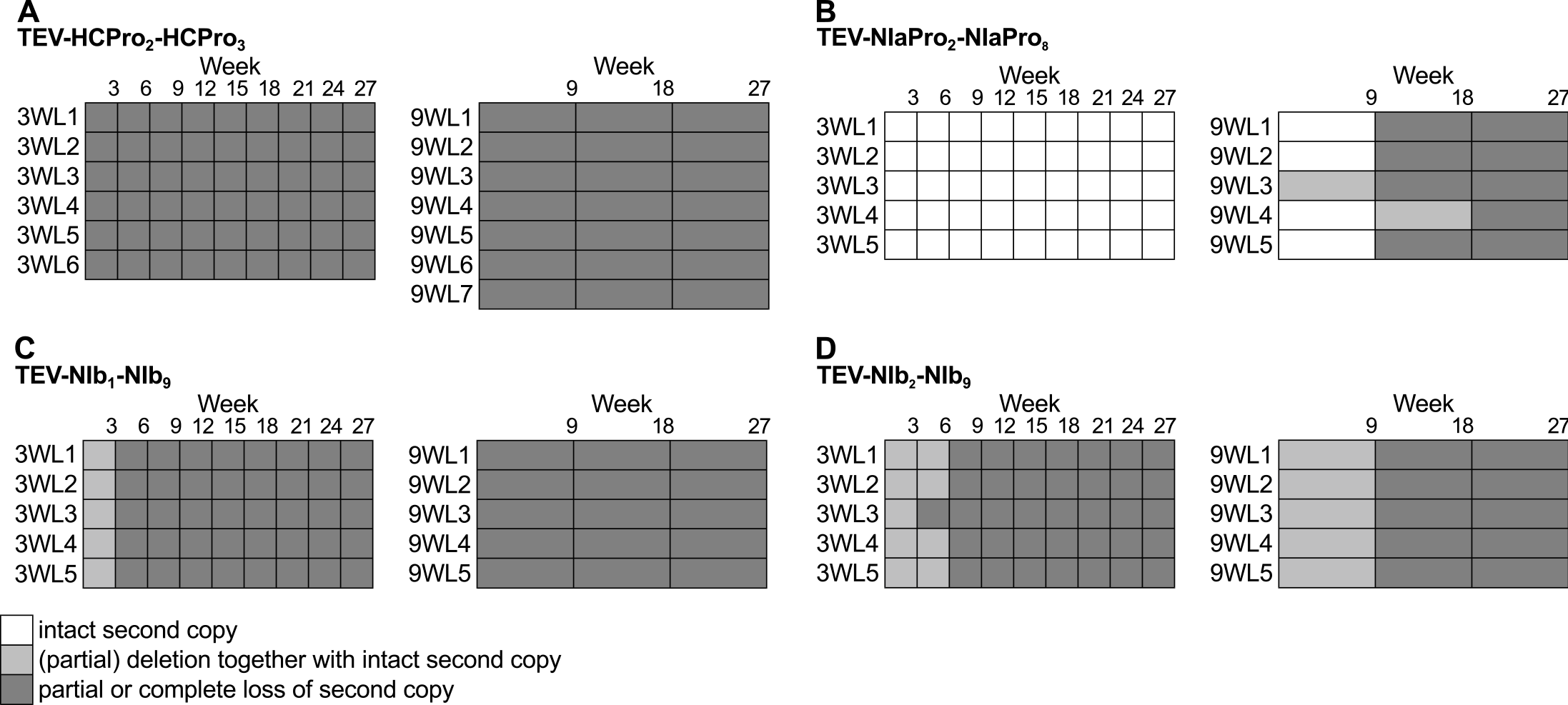
Deletion detection along the evolution experiments. RT-PCR was performed on the region containing a duplication in the viral genotypes (A-D). Either an intact duplicated copy (white boxes), a deletion together with an intact duplicated copy (light-grey boxes), or a partial or complete loss of the duplicated copy (dark-grey boxes) were detected.

**Figure 3.**
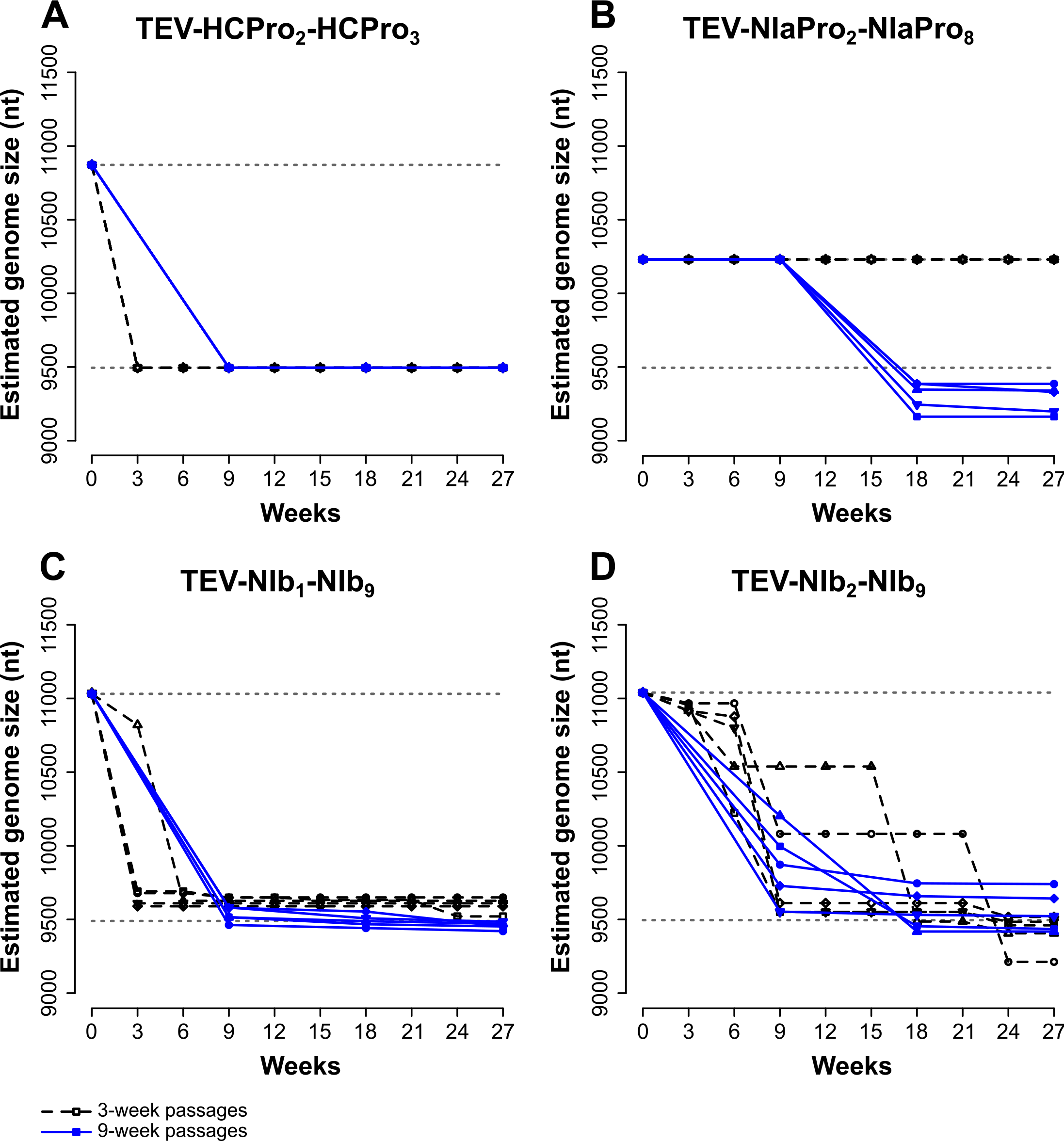
The reduction in genome size over time. The different panels display how the genome size of the different viral genotypes with gene duplications (A-D) changes along the evolution experiments. The dotted grey lines indicate the genome sizes of the wild-type virus (below) and the ancestral viruses (above). The genome sizes of the 3-week lineages are drawn with dashed black lines and open symbols, and those of the 9-week lineages are drawn with continuous blue lines and filled symbols.

### Viruses with a gene duplication have reduced fitness which cannot always be fully restored after deletion

For both the ancestral and evolved virus populations, we measured within-host competitive fitness (Figure 4) and viral accumulation (Figure 5). Comparing the ancestral viruses containing a gene duplication to the ancestral wild-type virus (solid circles in Figures 4 and 5), we observed statistically significant decreases in competitive fitness (Figure 4A; TEV-HCPro_2_-HCPro_3_: *U* = 8.398, *P* = 0.001. Figure 4B; TEV-NIaPro_2_-NIaPro8: *U* = 12.776, *P* < 0.001. Figure 4C; TEV-NIbrNIb_9_: *h* = 6.379, *P* = 0.003; TEV-NIb_2_-NIb_9_: *U* = 8.348, *P* = 0.001). Statistically significant decreases in accumulation levels were also observed for TEV-HCPro_2_-HCPro_3_, TEV-NIb1-NIb_9_ and TEV-NIb_2_-NIb9 (Figure 5A; TEV-HCPro_2_-HCPros: *t_A_* = 3.491, *P* = 0.0251. Figure 5C; TEV-NIbrNIb9: *t_A_* = 45.097, *P* < 0.001; TEV-NIb_2_-NIb_9_: t_4_ = 8.650, *P* < 0.001). However, there was no difference in accumulation for the virus with a duplication of *NIa-Pro* (Figure 5B; TEV-NIaPro_2_-NIaPro_8_: t_4_ = 2.099, *P* = 0.104). All the virus with duplications therefore have a reduced within-host competitive fitness, and three out of four viruses also have reduced accumulation. None the possible benefits of these gene duplications therefore can compensate for their costs.

**Figure 4.**
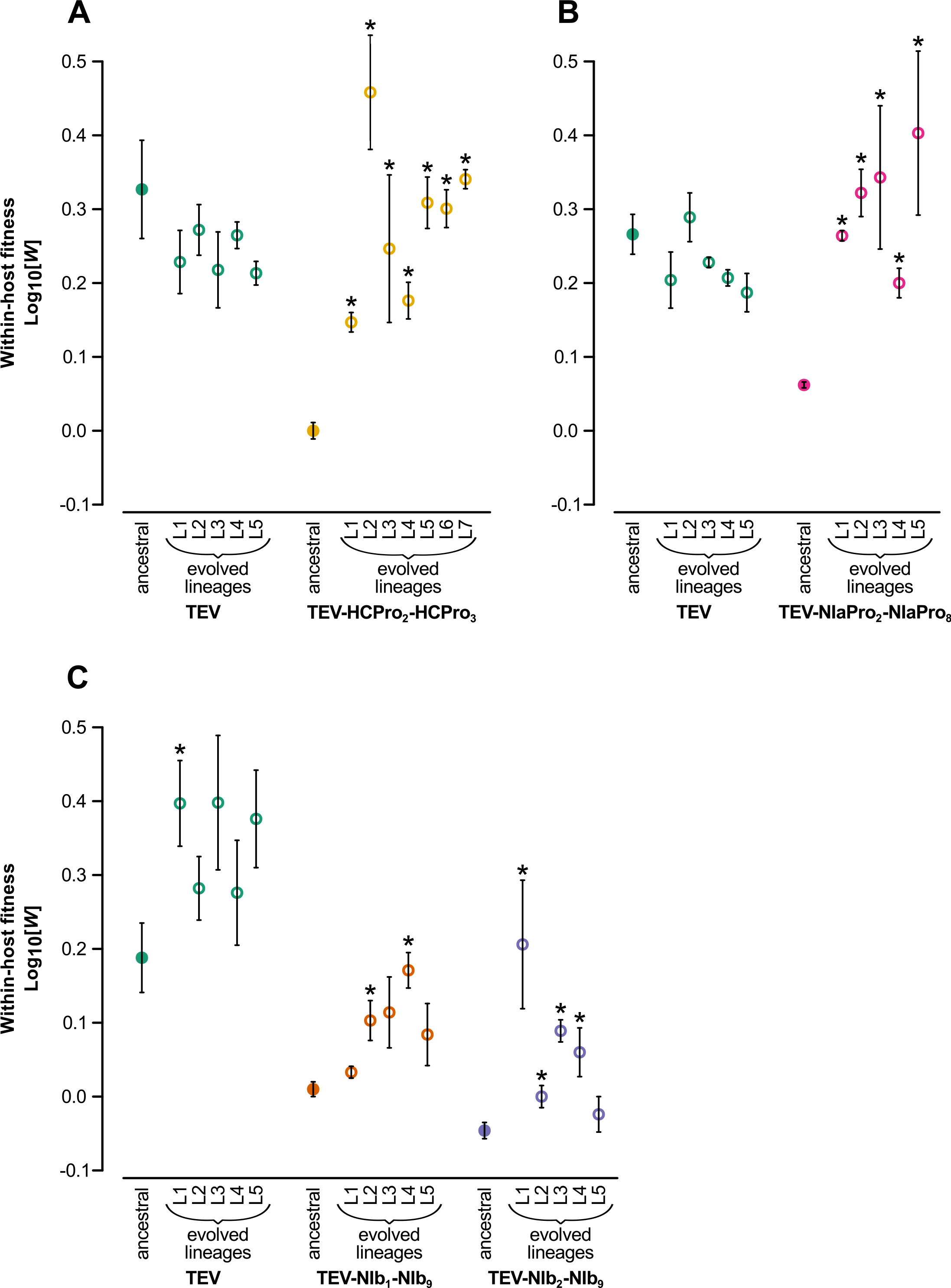
Within-host competitive fitness of the evolved and ancestral lineages. Fitness (*W*), as determined by competition experiments and RT-qPCR of the different viral genotypes with respect to a common competitor; TEV-eGFP. The ancestral lineages are indicated by filled circles and the evolved lineages by open circles. The different viral genotypes are color coded, where the wild-type virus is drawn in green. The asterisks indicate statistical significant differences of the evolved lineages as compared to their corresponding ancestral lineages (*t*-test with Holm-Bonferroni correction).

**Figure 5.**
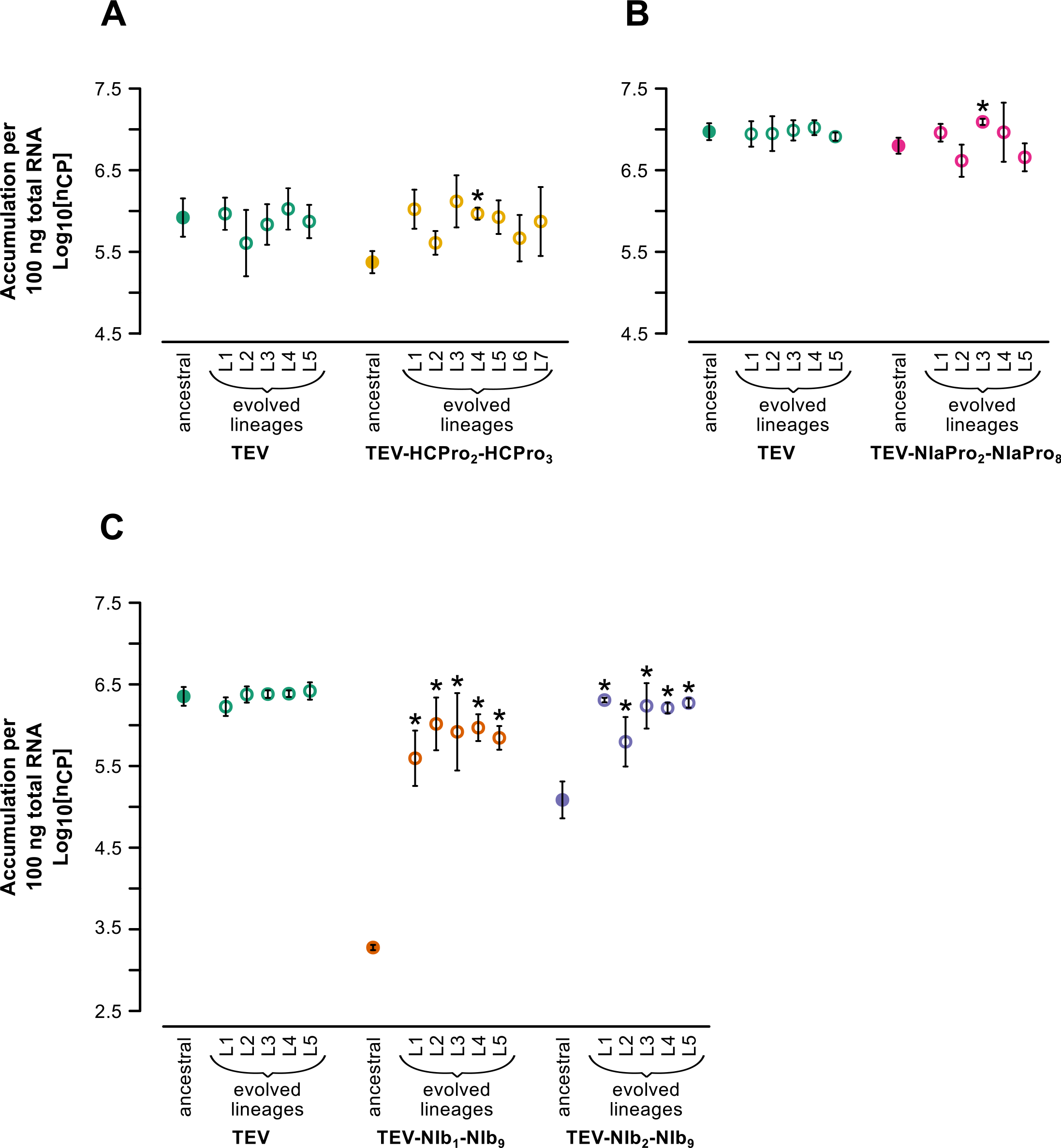
Virus accumulation of the evolved and ancestral lineages. Virus accumulation, as determined by accumulation experiments and RT-qPCR at 7 dpi of the different viral genotypes. The ancestral lineages are indicated by filled circles and the evolved lineages by open circles. The different viral genotypes are color coded, where the wild-type virus is drawn in green. The asterisks indicate statistical significant differences of the evolved lineages as compared to their corresponding ancestral lineages (*t*-test with Holm-Bonferroni correction).

After evolving the viruses with gene duplications using three 9-week passages, within-host competitive fitness of all four viruses increased (open circles in Figure 4), compared to their respective ancestral viruses (Figure 4; asterisks indicate a significant increase, *t*-test with Holm-Bonferroni correction). Within-host fitness was similar to the evolved wild-type TEV for evolved lineages of both TEV-HCPro_2_-HCPro_3_ (Figure 4A; Mann-Whitney *U* = 23, *P* = 0.432) and TEV-NIaPro_2_-NIaPro_8_ (Figure 4B; Mann-Whitney *U* = 20, *P* = 0.151). On the other hand, the evolved TEV-NIb_1_-NIb_9_ and TEV-NIb_2_-NIb_9_ lineages did not reach wild-type virus within-host fitness levels (Figure 4C; TEV-NIb_1_-NIb_9_: Mann-Whitney *U* = 0, *P* = 0.008; TEV-NIb_2_-NIb_9_: Mann-Whitney *U* = 0, *P* = 0.008). The within-host fitness of evolved lineages was also compared by means of a nested ANOVA (Table 2), allowing the independent lineages (at least 5) to be nested within the genotype and the independent plant replicates (3) to be nested within the independent lineages within the genotype. The nested ANOVA confirms that there is indeed an effect of the genotype for the TEV-NIb_1_-NIb_9_ and TEV-NIb_2_-NIb_9_ viruses (Table 2 and Willemsen et al. 2016), while for the TEV-HCPro_2_-HCPro_3_ and TEV-NIaPro_2_-NIaPro_8_ no effect was found. In summary, the fitness of TEV-HCPro_2_-HCPro_3_ and TEV-NIaPro_2_-NIaPro_8_ clearly increases to levels similar to the wild-type, whilst fitness did not increase for TEV-NIb_2_-NIb_9_ and TEV-NIb_1_-NIb_9_.

Together with within-host fitness, virus accumulation also increased significantly for the evolved TEV-NIb_1_-NIb_9_ and TEV-NIb_2_-NIb_9_ virus lineages (Figure 5C; asterisks), when compared to their respective ancestral viruses. However, accumulation levels did not increase significantly for most of the evolved lineages of the TEV-HCPro_2_-HCPro_3_ and TEV-NIaPro_2_-NIaPro_8_ genotypes. Nevertheless, these two genotypes have much higher initial accumulation levels than the genotypes with a duplication of the *NIb* gene. When comparing the accumulation levels of the evolved lineages to those of the wild-type (again, using lineage as the replication unit), TEV-HCPro_2_-HCPro_3_ (Figure 5A; Mann-Whitney *U* = 20, *P* = 0.755), TEV-NIaPro_2_-NIaPro8 (Figure 5B; Mann-Whitney *U* = 11, *P* = 0.841), and TEV-NIb_2_-NIb_9_ (Figure 5C; Mann-Whitney *U* = 3, *P* = 0.056) do reach wild-type accumulation levels, whilst TEV-NIb_1_-NIb_9_ does not (Figure 5C; Mann-Whitney *U* = 0, *P* = 0.008). Comparing the accumulation levels of the evolved lineages by means of a nested ANOVA (Table 2) confirms that there is an effect of the genotype for the TEV-NIb_1_-NIb_9_ virus (Table 2 and Willemsen et al. 2016), while no effect for the TEV-HCPro_2_-HCPro_3_, TEV-NIaPro_2_-NIaPro_8_ and TEV-NIb2-NIb9 viruses was found.

**Table 2.** Nested ANOVA s on within-host competitive fitness and viral accumulation

| Genotype | Trait | Source of Variation | SS | df | MS | *F* | *P* |
| --- | --- | --- | --- | --- | --- | --- | --- |
| TEV-HCPro<sub>2</sub>-<br>HCPro<sub>3</sub> | Competitive fitness | Genotype | 0.049 | 1 | 0.049 | 0.786 | 0.396 |
|  |  | Lineage within Genotype | 0.624 | 10 | 0.062 | 9.947 | < 0.001 |
|  |  | Plant within Lineage within Genotype | 0.150 | 24 | 0.006 | 347.244 | < 0.001 |
|  |  | Error | 0.001 | 72 | 1.81×10<sup>-5</sup> |  |  |
|  | Accumulation | Genotype | 0.008 | 1 | 0.008 | 0.080 | 0.783 |
|  |  | Lineage within Genotype | 1.012 | 10 | 0.101 | 1.005 | 0.467 |
|  |  | Plant within Lineage within Genotype | 2.417 | 24 | 0.101 | 51.187 | < 0.001 |
|  |  | Error | 0.142 | 72 | 0.002 |  |  |
| TEV-NIaPro<sub>2</sub>-<br>NIaPro<sub>8</sub> | Competitive fitness | Genotype | 0.155 | 1 | 0.155 | 4.534 | 0.066 |
|  |  | Lineage within Genotype | 0.274 | 8 | 0.034 | 4.285 | 0.004 |
|  |  | Plant within Lineage within Genotype | 0.160 | 20 | 0.008 | 503.714 | < 0.001 |
|  |  | Error | 0.001 | 60 | 1.59×10<sup>-5</sup> |  |  |
|  | Accumulation | Genotype | 0.246 | 1 | 0.246 | 1.199 | 0.305 |
|  |  | Lineage within Genotype | 1.643 | 8 | 0.205 | 2.224 | 0.070 |
|  |  | Plant within Lineage within Genotype | 1.847 | 20 | 0.092 | 323.631 | < 0.001 |
|  |  | Error | 0.017 | 60 | 0.001 |  |  |
| TEV-NIb<sub>1</sub>-<br>NIb<sub>9</sub> | Competitive fitness | Genotype | 1.308 | 1 | 1.308 | 49.734 | < 0.001 |
|  |  | Lineage within Genotype | 0.212 | 8 | 0.026 | 3.145 | 0.018 |
|  |  | Plant within Lineage within Genotype | 0.169 | 20 | 0.008 | 175.319 | < 0.001 |
|  |  | Error | 0.003 | 58 | 4.81×10<sup>-5</sup> |  |  |
|  | Accumulation | Genotype | 5.374 | 1 | 5.374 | 36.006 | < 0.001 |
|  |  | Lineage within Genotype | 1.194 | 8 | 0.149 | 0.939 | 0.507 |
|  |  | Plant within Lineage within Genotype | 3.178 | 20 | 0.159 | 263.098 | < 0.001 |
|  |  | Error | 0.036 | 60 | 0.001 |  |  |
| TEV-NIb<sub>2</sub>-<br>NIb<sub>9</sub> | Competitive fitness | Genotype | 1.796 | 1 | 1.796 | 36.175 | < 0.001 |
|  |  | Lineage within Genotype | 0.397 | 8 | 0.050 | 4.207 | 0.004 |
|  |  | Plant within Lineage within Genotype | 0.236 | 20 | 0.012 | 519.611 | < 0.001 |
|  |  | Error | 0.001 | 60 | 2.27×10<sup>-5</sup> |  |  |
|  | Accumulation | Genotype | 0.824 | 1 | 0.824 | 3.728 | 0.090 |
|  |  | Lineage within Genotype | 1.771 | 8 | 0.221 | 3.439 | 0.012 |
|  |  | Plant within Lineage within Genotype | 1.290 | 20 | 0.065 | 110.478 | < 0.001 |
|  |  | Error | 0.034 | 59 | 0.001 |  |  |

When comparing the within-host competitive fitness of the evolved TEV-NIaPro2-NIaPro8 9-week lineages to the 3-week lineages, we found that there is a linear relationship between genome size and within-host competitive fitness (Figure 6; Spearman’s rank correlation *ρ* = −0.795, 10 d.f., *P* = 0.006). The evolved 9-week lineages, that contain genomic deletions, have a significant higher within-host competitive fitness (Mann-Whitney *U* = 4.5, *P* < 0.001) than the evolved 3-week lineages without deletions.

**Figure 6.**
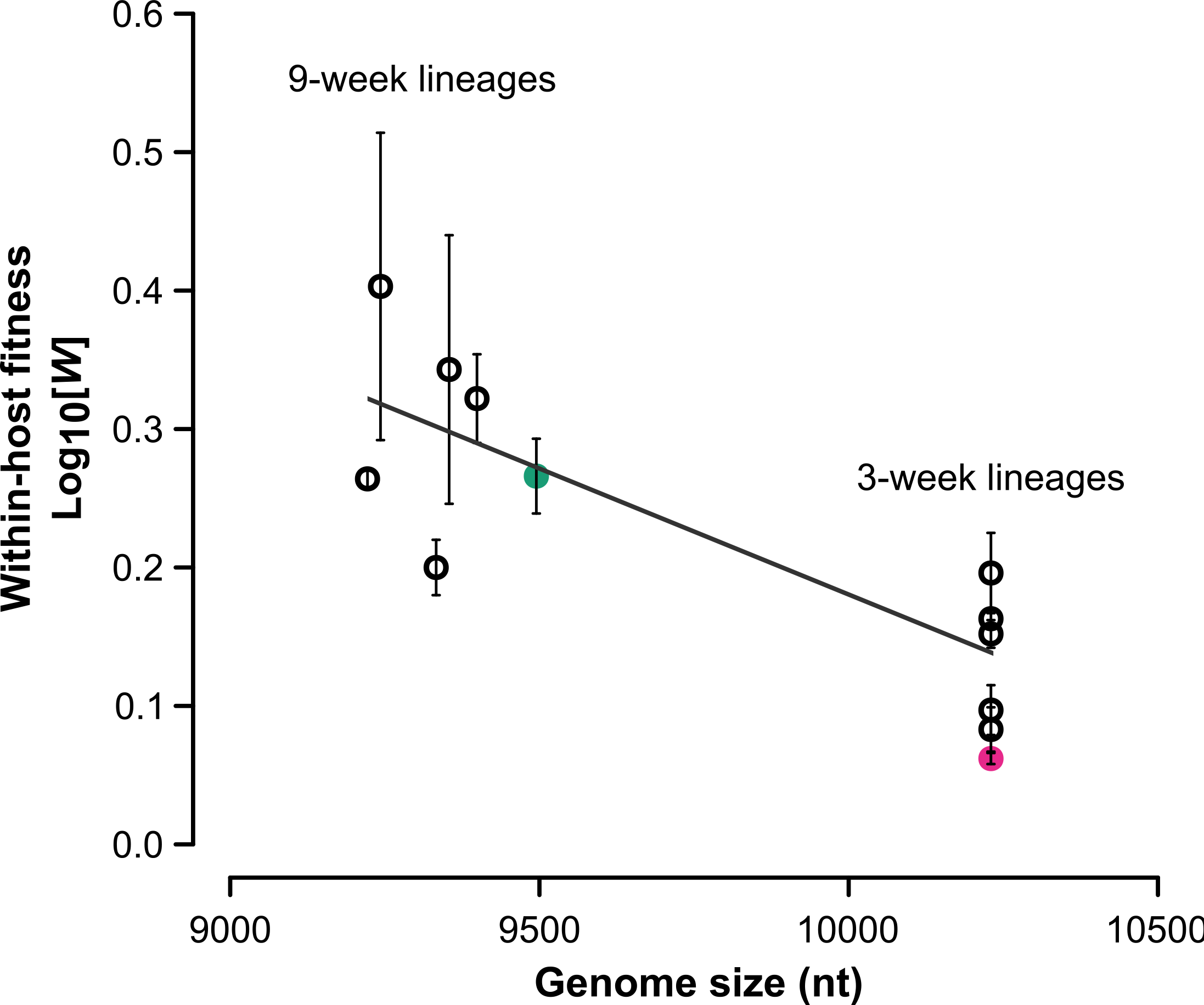
The relationship between genome size and within-host competitive fitness. The pink filled circle represents the within-host competitive fitness of the ancestral TEV-NIaPro_2_-NIaPro_8_ and the green filled circle that of the ancestral wild-type TEV. The black open circles represent the evolved 3-week (right) and 9-week (left) TEV-NIaPro_2_-NIaPro_8_ lineages. The evolved 9-week lineages, that contain genomic deletions, have a significant higher within-host competitive fitness (Mann-Whitney *U* = 4.5, *P* < 0.001) than the evolved 3-week lineages without deletions. A linear regression has been drawn to emphasize the trend in the data.

### Genome sequences of the evolved lineages

All evolved and ancestral lineages have been fully sequenced using the Illumina technology. The sequences of the ancestral lineages were used as an initial reference for the evolved lineages. Furthermore, for the lineages where deletions were detected by RT-PCR (Figure 3), parts of the genome were sequenced by Sanger to determine the exact deletion sites. The majority deletions variants were used to construct new reference sequences for each of the evolved TEV-HCPro_2_-HCPro_3_, TEV-NIaPro_2_-NIaPro_8_, TEV-NIb_r_NIb9, and TEV-NIb_2_-NIb9 lineages that contain deletions. After an initial mapping step, mutations were detected in the evolved lineages as compared to their corresponding ancestor (Materials and Methods).

Beside the large genomic deletions, different patterns of adaptive evolution were observed for each viral genotype (Figure 7 and Table 3). For the evolved TEV-HCPro_2_-HCPro_3_ virus a convergent nonsynonymous mutation was found in 3/7 9-week lineages in the *P1* gene (A304G), however, this mutation was also present in 1/5 9-week lineages of TEV. Another convergent nonsynonymous mutation was found in 3/7 9-week lineages in the *P3* gene (U4444C), known to be implicated in virus amplification and host adaptation (Revers & Gartia 2015). For the evolved TEV-NIaPro_2_-NIaPro_8_ virus, fixed convergent nonsynonymous mutations were found in the duplicated *NIa-Pro* (C1466U) copy in 4/5 3-week lineages, and in *6K1* (A4357G) in 3/5 3-week lineages. The latter mutation was also fixed in 1/5 3-week TEV lineages. For the evolved TEV-NIb_1_-NIb_9_ virus a fixed convergent nonsynonymous mutation was found in the pseudogenized *NIb* copy (A1643U) in 2/5 3-week lineages. For the evolved TEV-NIb_2_-NIb_9_ virus one fixed synonymous mutation was found in the multifunctional CI protein (C6531U) in 2/5 9-week lineages. Other convergent mutations in all virus genotypes were found in *VPg*, *NIa-Pro* and *NIb* genes, however these mutations were also found in 2 or more lineages of the wild-type virus (Figure 7). Therefore, we do not consider these genotype-specific mutations as adaptive.

**Table 3.** Adaptive convergent mutations within each virus genotype

| **Virus genotype** | **nt change at**<br>**ancestral position** | **aa**<br>**change** | **gene** | **nt position**<br>**in gene** |
| --- | --- | --- | --- | --- |
| TEV-HCPro<sub>2</sub>-HCPro<sub>3</sub> | A304G | I→V | P1 | 160 |
|  | U4444C | S→P | P3 | 625 |
| TEV-NIaPro<sub>2</sub>-NIaPro<sub>8</sub> | C1466U | S→L | NIa-Pro<sub>2</sub> | 410 |
|  | A4357G | I→V | 6K1 | 148 |
| TEV-NIb<sub>1</sub>-NIb<sub>9</sub> | A1643U | Y→F | NIb<sub>1</sub> | 1499 |
| TEV-NIb<sub>2</sub>-NIb<sub>9</sub> | C6351U | Y→Y | CI | 1173 |

nt: nucleotide; aa: amino acid

**Figure 7.**
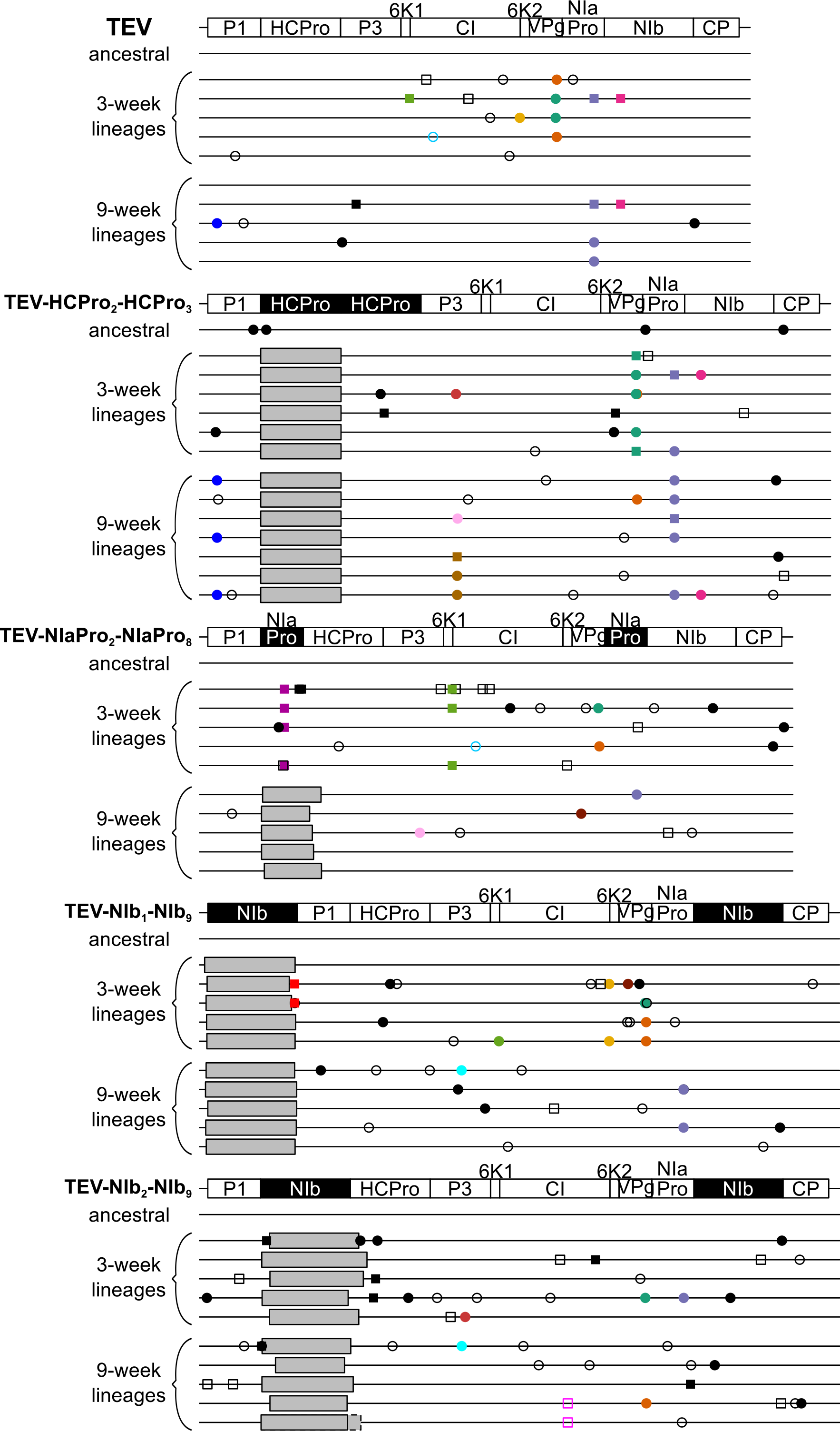
The relationship between genome size and within-host competitive fitness. Mutations were detected using NGS data of the evolved virus lineages as compared to their ancestral lineages. The square symbols represent mutations that are fixed (> 50%) and the circle symbols represent mutations that are not fixed (< 50%). Filled symbols represent nonsynonymous substitutions and open symbols represent synonymous substitutions. Black substitutions occur only in one lineage, whereas color-coded substitutions are repeated in two or more lineages, or in a lineage from another virus genotype. Note that the mutations are present at different frequencies as reported by SAMtools. Grey boxes indicate genomic deletions in the majority variant.

After remapping the cleaned reads against a new defined consensus sequence for each lineage, we looked at the variation within each lineage. Single nucleotide polymorphisms (SNPs) were detected at a frequency as low as 1%. In the evolved TEV-HCPro_2_-HCPro_3_ lineages, a total of 633 SNPs were detected, with a median of 45 (range 27-247) per lineage. There is no clear difference in the number of SNPs comparing the evolved 3-week lineages, median 48.50 (35-247), and the evolved 9-week lineages, median 44 (27-56). However, there is one 3-week lineage (3WL6) that accumulated a much higher number of SNPs (247) compared to the other TEV-HCPro2-HCPro3 lineages. In the evolved TEV-NIaPro2-NIaPro8 lineages, a total of 421 SNPs were detected, with a median of 43.50 (11-103) per lineage. The evolved 3-week lineages that have not lost the duplicated *NIa-Pro* copy accumulated more SNPs per lineage, median 53 (40-59), compared to the evolved 9-week lineages, median 33 (11-103). However, one lineage of the 9-week lineages (9WL2) also accumulated a much higher number of SNPs (103) compared to the other TEV-NIaPro_2_-NIaPro_8_ lineages. In the evolved wild-type TEV lineages, a total number of 402 SNPs were detected, with a median of 35 (17-63) SNPs per lineage. Like for TEV-HCPro_2_-HCPro_3_, for TEV there is no clear difference in the number of SNPs comparing the evolved 3-week lineages, median 34 (32-50), and the evolved 9-week lineages, median 36 (17-63). The data for the TEV-NIb_1_-NIb_9_ and TEV-NIb_2_-NIb_9_ lineages can be found in Willemsen et al. (2016), where a total 301 and 220 SNPs were detected with medians of 36 (27-45) and 23.5 (4-44) per lineage, respectively. In all four virus genotypes as well as in the wild-type virus, most of the SNPs were present at low frequency (SNPs < 0.1: TEV-HCPro_2_-HCPro_3_ = 89.8%; TEV-NIaPro_2_-NIaPro_8_ = 84.8%; TEV-NIb1-NIb9 = 83.3%; TEV-NIb_2_-NIb9 = 81.2%; TEV = 85.2%), with a higher prevalence of synonymous (TEV-HCPro_2_-HCPro_3_: 54.7%, TEV-NIaPro_2_-NIaPro8: 57.7%, TEV-NIb_1_-NIb_9_: 66.4%, TEV-NIb_2_-NIb_9_: 64.5%, TEV: 59.7%) versus nonsynonymous changes (supplementary Figure S1, Supplementary Material online). Moreover, a percentage of the nonsynonymous changes for TEV-HCPro_2_-HCPro_3_ (16.7%) and TEV-NIaPro_2_-NIaPro_8_ (7.3%) as well as the wild-type TEV (14.8%), are actually leading to stop codons and therefore unviable virus variants. For both TEV-HCPro_2_-HCPro3 and TEV-NIaPro_2_-NIaPro_8_ the difference in the distribution of synonymous versus nonsynonymous SNP frequency is significant (Kolmogorov-Smirnov test; TEV-HCPro_2_-HCPros: *D* = 0.219, *P* < 0.001; TEV-NIaPro_2_-NIaPro_8_: *D* = 0.151, *P* = 0.009), whilst for TEV-NIb_1_-NIb_9_ and TEV-NIb_2_-NIb_9_ (Willemsen et al. 2016) and the wild-type virus this is not significant (Kolmogorov-Smirnov test; TEV: *D* = 0.084, *P* = 0.555). For more details on the frequency of the SNPs within every lineage, see supplementary Tables S3, S4 and S5 (Supplementary Material online).

### Genomic stability of TEV with duplications of homologous genes

To better understand the evolutionary dynamics of viruses with gene duplications, we developed a simple mathematical model of virus competition and evolution. Based on amplicon sizes, the genome size for all evolved lineages was estimated for every passage (Figure 3). Our model attempts to account for these data, and specifically how long the duplicate gene copy is maintained in the virus population. The model we developed describes how a population composed initially of only virus variants with a gene duplication (variant *A*) through recombination, selection and genetic drift acquires and eventually fixes a new variant that only retains the original copy of the duplicated gene (variant *B*). The model includes a genetic bottleneck at the start of each round of passaging (*i.e.*, the initiation of infection in the inoculated leaf), with a fixed total number of founders and binomially distributed number of founders for variants containing the gene duplication. Following this genetic bottleneck, there is deterministic growth of both variants as well as deterministic recombination of *A* into *B*. The main question we addressed is whether knowing the fitness of duplicated viruses (*i.e., a*) is sufficient information to predict the stability of the inserted gene. Or do the data support a context-dependent recombination rate, with the context being (*i*) identity and position of the duplication, (*ii*) passage length, or (*iii*) both? We considered four different situations that are represented in the following models:

Model 1: one recombination rate for all conditions (1 parameter);

Model 2: virus-genotype-dependent recombination rate (4 parameters);

Model 3: passage-duration-dependent recombination rates (2 parameters);

Model 4: virus-genotype-and passage-duration-dependent recombination rates (full model, 8 parameters).

The model estimates of *δ* are given in Table 4. Note that the parameter is often a minimum (when the virus is very unstable) or a maximum value (when the virus is very stable). If the optimum is represented by more than one parameter value, the mean of these values is given. When comparing these models, we found that Model 2 is the best-supported model (Table 5). Thus, only a genotype-dependent recombination rate is required to account for the data. The results strongly suggest that knowing the fitness of a virus with a gene duplication is not sufficient information for predicting genomic stability. Rather, as the recombination rate is dependent on the genetic context, the supply of recombinants which have lost the duplicated gene will vary greatly from one genotype to another. High recombinogenic sites will remove the second copy fast, while low recombinogenic sites will preserve the copy for longer periods of time, after which it will be unavoidably removed, as confirmed by the numerical analysis of equations 1 and 2 (supplementary Text S1, Supplementary Material online).

**Table 4.**
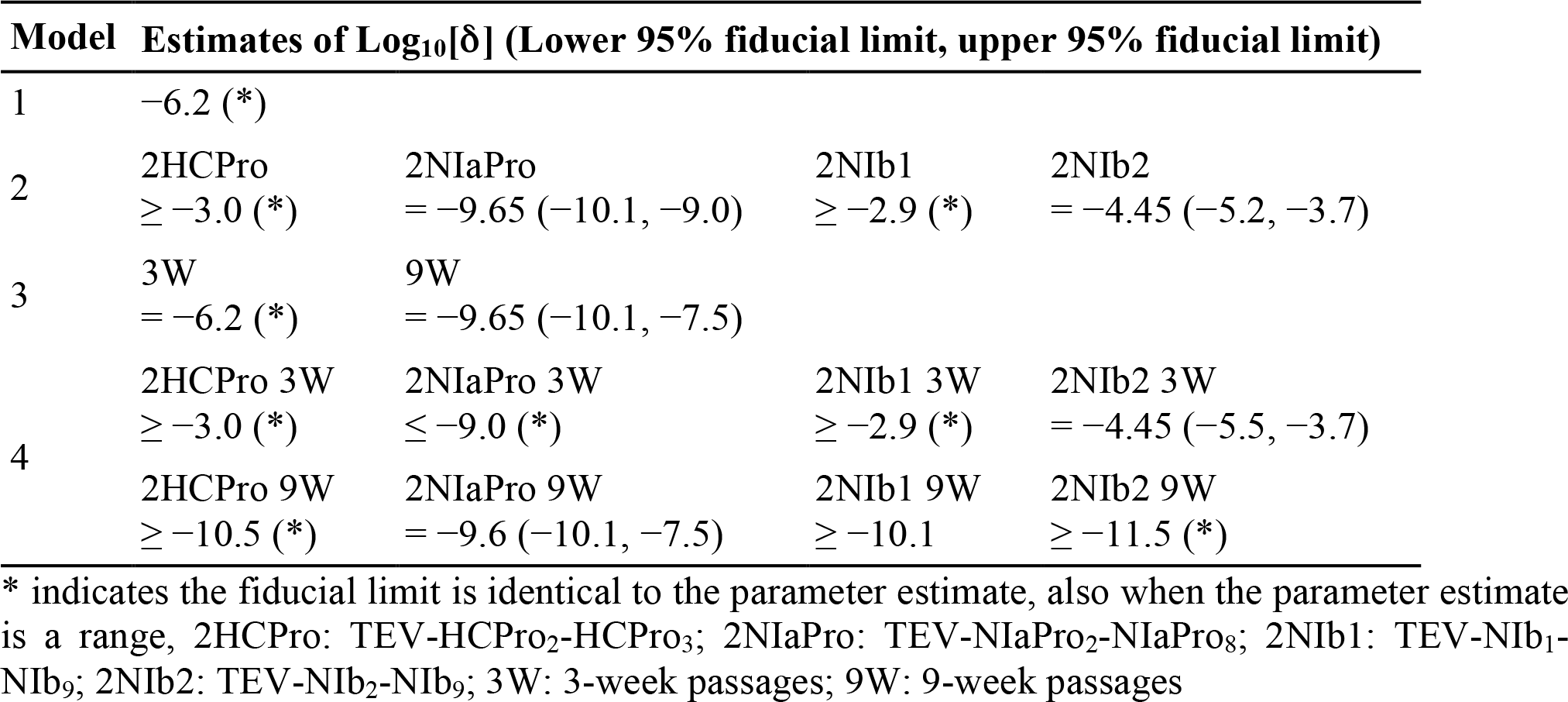
Model parameter estimates for deterministic recombination rate

**Table 5.** Model selection for models with deterministic recombination

| **Model** | **Parameters** | **NLL** | **AIC** | **ΔAIC** | **Akaike Weight** |
| --- | --- | --- | --- | --- | --- |
| 1 | 1 | 254.284 | 510.569 | 443.470 | 0.000 |
| 2 | 4 | 29.549 | 67.099 | −0.000 | 0.982 |
| 3 | 2 | 226.669 | 457.339 | 390.240 | 0.000 |
| 4 | 8 | 29.549 | 75.099 | 8.000 | 0.018 |

On the other hand, passage duration has a strong effect on the observed stability of gene duplications. However, model selection shows that this phenomenon can be sufficiently explained by considering the combined effects of selection and genetic drift, and without invoking passage-duration-dependent recombination rates. Given that recombination and selection are deterministic in the model, deletion variants will always arise during infection. However, depending on the rates of recombination and selection, these deletion variants may not reach a high enough frequency to ensure they are sampled during the genetic bottleneck at the start of each round of infection. This effect will be much stronger in the 3-week passages-since there will be fewer recombinants and less time for selection to increase their frequency-explaining why for some viruses there is such a marked difference in the observed genomic stability for different passage lengths.

The deterministic dynamics of the evolutionary model of stability of genomes containing gene duplications have also been investigated (supplementary Text S1, Supplementary Material online). These analyses were performed on the model as described in equations 1 and 2, albeit with a time-independent carrying capacity. The stability of three fixed points was analyzed: (i) the extinction of both *A* and B, (*ii)* the domination of *B* over the population, and (iii) the coexistence of *A* and *B* in the population. The fixed points analysis indicates that when *a* < *b* and *δ > a-b*, the *B* virus subpopulation will outcompete the *A* subpopulation. This parametric combination is the one obtained using the biologically meaningful parameter values shown in Table 1. This means that, under the model assumptions, the population of virions containing a gene duplication is unstable, and the population will be asymptotically dominated by virions containing a single gene copy. Notice that, the rate of recombination (δ) is also involved in this process of outcompetition (see the bifurcation values calculated in supplementary Text S1, Supplementary Material online). Specifically, if δ > *a-b*, coexistence of the two virion types is not possible. Interestingly, the model also reveals the existence of a transcritical bifurcation separating the scenario between coexistence of the two types of virions and dominance of *B*. Such a bifurcation can be achieved by tuning *δ* as well as by unbalancing the fitness of *A* and *B* virus types. This bifurcation gives place to a smooth transition between these two possible evolutionary asymptotic states (*i.e.*, unstable *A* population and coexistence of *A* and *B* populations). The bifurcation will take place when *a = b + δ*, or when *δ* is above a critical threshold, *δ_c_ = a-b*, that can be calculated from the mathematical model (supplementary Text S1, Supplementary Material online).

Finally, we characterized the time to extinction of the *A* subpopulation. We characterized these extinction times as a function of viral fitness and of δ. Such a time is found to increase super-exponentially as the fitness of the *A* subpopulation, approaches the fitness of the *B* subpopulation. This effect is found for low values of recombination rates, similar to those shown in Table 4. As expected, increases in *δ* produce a drastic decrease in the time needed for the single-copy population to outcompete the population of viruses with the duplicated genes (supplementary Text S1, Supplementary Material online).

## Discussion

Genetic redundancy is thought to be evolutionary unstable in viruses due to the costs associated with maintaining multiple gene copies. Here we tested the stability of duplicated sequences that might contribute to the enhancement of a virus function or even exploration of new functions. Overall, none of the duplication events explored appeared to be beneficial for TEV, both in terms of their immediate effects and second-order effects on evolvability. Gene duplication resulted in either an inviable virus or a significant reduction in viral fitness. In all cases the duplicated gene copy, rather than the ancestral gene copy, was deleted during long-duration passages. The earlier detection and more rapid fixation of deletion variants during longer-duration passages is congruent with results from a previous study (Zwart et al. 2014), where deletions of *eGFP* marker inserted in the TEV genome were usually observed after a single 9-week passage, but were rarely spotted even after nine 3-week passages. On the other hand, the highly diverging results for genome stability obtained here for the different viruses, suggests that passage duration is not the main factor determining whether gene duplication will be stable. Therefore, the size of the duplicated gene, the nature of the gene, and/or the position for duplication could play a role in the stability of genomes with a duplication.

The observation that gene duplication events result in decreases in fitness is not unexpected. The high mutation rate of RNA viruses is likely to constrain genome size (Holmes 2003), given that most mutations are deleterious (Sanjuán et al. 2004; Elena et al. 2006). This imposes the evolution of genome compression, where overlapping reading frames play a major role (Belshaw et al. 2007; Belshaw et al. 2008). In our model virus we speculate that the fitness cost can be related to three processes: (*i*) the increase in genome size, (*ii*) the extra cost of more proteins being expressed in the context of using more cellular resources, and (iii) a disturbance in correct polyprotein processing. Although all the duplications considered here could conceivably have advantages for viral replication or encapsidation, our results suggest that the any such advantages are far outweighed by the costs associated with a larger genome, increased protein expression or the effects on polyprotein processing. However, the duplication of *NIa-Pro* does not affect the viral accumulation rate. This could be explained by the fact that the *NIa-Pro* gene is much smaller than the other duplicated genes. Consequently, in conditions where selection has the least time to act between bottleneck events associated with infection of a new host (3-week passages), no deletions were observed. In the long-passage experiment selection has more time to act and increase the frequency of beneficial *de novo* variants, allowing them to be sampled during the bottleneck at the start of the next round of infection. In addition, the size of the gene duplication also seems to play a role. But what about the position and the nature of the duplicated gene? When duplicating the same gene, *NIb*, to either the first or second position in the TEV genome, we see clear differences in the deletion dynamics and fitness measurements (Figures 3, 4, 5 and Willemsen et al. 2016). Comparing the duplication and subsequent deletion of *HC-Pro*, *NIa-Pro* and *NIb* at the same second position, we observe that both accumulation and within-host competitive fitness cannot be completely restored by the virus that originally had two copies of the *NIb* replicase gene, whilst viruses that originally had two copies of the *HC-Pro* gene or the main viral protease *NIa-Pro* gene do restore their fitness after deletion. However, since our evolutionary experiments were limited to approximately half a year, we cannot rule out complete restoration of fitness over longer time periods.

At the sequence level, there were some convergent single-nucleotide mutations, although in most cases these occur only in a small fraction of lineages. The transient presence of the duplicated *NIa-Pro* copy in the 3-week lineages does seem to be linked to an adaptive mutation. However, our fitness measurements suggest that the cost of gene duplication cannot be overcome by this single nucleotide mutation. The main change in the evolved lineages is the deletion of the duplicated gene copy. However, some deletions extend beyond the duplicated gene copy, including the N-terminal region of the HC-Pro cysteine protease, similar to results obtained by previous studies (Dolja et al. 1993; Zwart et al. 2014; Willemsen et al. 2016). The N-terminal region of HC-Pro is implicated in transmission by aphids (Thornbury et al. 1990; Atreya et al. 1992) and is not essential for viral replication and movement (Dolja et al. 1993; Cronin et al. 1995). Our experimental setup does not involve transmission by aphids, however, we do not observe this deletion when evolving the wild-type virus. Moreover, we only observe this deletion when the position of gene duplication or insertion (Zwart et al. 2014) is before HC-Pro, suggesting that gene insertion and subsequent deletion at this position facilitates recombination to an even smaller genome size.

By fitting a mathematical model of virus evolution to the data, we find that knowing only fitness is not enough information to predict the stability of the duplicated genes: as we have shown, there is a context-dependent recombination rate, and specifically, the identity and position of the duplication also play a role. Given that the supply rate of variants with large deletions will be driven largely by homologous recombination, we had expected stability to depend on the genetic context. The estimates of the recombination rate per genome and generation in this study are far lower than previously reported for TEV, which was estimated to be 3.427×10^−5^ per nucleotide and generation (Tromas et al. 2014b), which translates into 0.3269 per genome and generation. The estimates of this study (TEV-HCPro_2_-HCPro_3_: 1.000×10-^3^; TEV-NIaPro_2_-NIaPro8: 2.239×10-^10^; TEV-NIb1-NIb9: 1.259×10-^3^; TEV-NIb_2_-NIb_9_: 3.548×10^−5^) are much closer to the per nucleotide estimate. This large discrepancy could be related to two factors. First, Tromas et al. (2014b) considered recombination between two highly similar genotypes, which requires consideration of many details of the experimental system, including the rate at which cells will be coinfected by these genotypes. On the other hand, considering these details should lead to a general estimate of the recombination rate (as opposed to the rate at which two different genotypes will recombine in a mixed infection) and hence this explanation is not very satisfactory. Second, only a small fraction all recombination events will render viruses with a conserved reading frame, and a suitable deletion size: large enough to have appreciable fitness gains and be selected, but small enough not to disrupt the surrounding cistrons or polyprotein processing. Therefore, the parameter *δ* described here is in fact the rate at which this particular subclass of recombinants occurs. This subclass is likely to be only a small fraction of all possible recombinants, and hence it is quite reasonable that these two estimates of the recombination rate vary by several orders of magnitude.

In addition to gene duplications, the model developed in this study can be applied to predict the stability of other types of sequence insertions, such as those brought about by horizontal gene transfer. Understanding the stability of gene insertions in genomes is highly relevant to the understanding genome-architecture evolution, but it also has important implications for biotechnological applications, such as heterologous expression systems. Our results here suggest that the fitness costs of extraneous sequences may not be a good predictor of genomic stability, in general. Therefore, in practical terms, it could be advisable to empirically test the stability of *e.g*. a viral construct, rather than make assumptions on stability based on parameters such as replication or accumulation.

## Acknowledgements

We thank Francisca de la Iglesia and Paula Agudo for excellent technical assistance.

## Supplementary files

### Supplementary file 1

This file contains supplementary Table S1 with primers flanking duplicated and ancestral gene regions from 5’ to 3’ and supplementary Table S2 with primers for Sanger sequencing from 5’ to 3’.

### Supplementary file 2

This file contains supplementary Figure S1 with the distribution of SNP frequencies in the evolved TEV-HCPro_2_-HCPro_3_, TEV-NIaPro_2_-NIaPro_8_ and TEV lineages.

### Supplementary file 3

This file contains supplementary Tables S3, S4 and S5 with the within population sequence variation of the evolved and ancestral TEV-HCPro_2_-HCPro_3_, TEV-NIaPro_2_-NIaPro_8_ and TEV lineages, respectively.

### Supplementary file 4

This file contains supplementary Text S1 where the numerical analysis of equations 1 and 2 is fully presented. The file also contains the figures referred in the supplementary text.

## Data Deposition

The sequences of the ancestral viral stocks were submitted to GenBank with accessions TEV-NIb_1_-NIb_9_: KT203712; TEV-NIb_2_-NIb_9_: KT203713; TEV-HCPro_2_-HCPro_3_: KX137150; TEV-NIaPro_2_-NIaPro_8_: KX137151; TEV: KX137149).

